# GPSNorm: Gaussian Process Spatial Normalization for Spatial Transcriptomics

**DOI:** 10.64898/2026.07.23.740319

**Authors:** Art Taychameekiatchai, Xiaowei Zhan, Guanghua Xiao, Peifeng Ruan

## Abstract

Spatial transcriptomics technologies enable measurement of gene expression while preserving spatial tissue organization, but they remain highly sensitive to technical variability such as library size differences, slide-level effects, and spatial artifacts. Most existing normalization approaches treat normalization as a preprocessing step and perform downstream analyses on normalized values as fixed inputs, ignoring the uncertainty introduced during normalization. We introduce **GPSNorm** (Gaussian Process Spatial Normalization), a Bayesian spatial normalization framework that jointly models technical variation, spatial structure, and biological signal within a unified hierarchical model. GPSNorm represents gene expression counts using a negative binomial latent Gaussian model whose spatial component is a Gaussian Markov random field approximating a Gaussian process and performs efficient approximate Bayesian inference using the Integrated Nested Laplace Approximation (INLA), producing posterior estimates that propagate normalization uncertainty into downstream differential expression analysis. In simulations anchored to empirical spatial transcriptomics data, GPSNorm accurately recovers spatial technical structure and improves log-fold change estimation compared with existing normalization methods. Applications to three spatial transcriptomics datasets—including human dorsolateral prefrontal cortex Visium data, a GeoMx COVID-19 lung damage study, and the Spatial Organ Atlas kidney dataset—demonstrate improved preservation of biologically expected spatial patterns and marker gene contrasts. These results show that jointly modeling normalization and downstream inference can improve the robustness and interpretability of spatial transcriptomics analyses.An open-source R implementation of GPSNorm is available at https://github.com/Tiny-Quant/GPSNorm.

## 1 Introduction

Spatial transcriptomics (ST) technologies enable genome-wide measurement of gene expression while preserving spatial coordinates within intact tissue sections. This makes it possible to study tissue architecture and spatially varying transcriptional programs that are not directly observable from dissociated sequencing data. (Ståhl et al.). Commercial platforms such as 10x Genomics Visium and NanoString GeoMx DSP have broadened the use of spatial profiling across biological and clinical applications. At the same time, observed counts in ST reflect both biological signal and technical variation, such as library size, slide-to-slide effects and within-slide spatial artifacts. Unlike in bulk and single-cell RNA sequencing settings, these technical components are spatially structured and make it difficult to distinguish unwanted technical variability from biologically meaningful signal. (Love et al.; Liu et al., a). For example, empirical studies have shown that library size can correlate with tissue architecture, creating potential confounding between technical and biological spatial structure (Bhuva et al.; Salim et al.).

This feature creates a statistical challenge for normalization. Most normalization approaches proceed sequentially: counts are first adjusted using global or spatially informed scaling factors, and downstream models are subsequently fit to the adjusted values. Methods originally developed for bulk and single-cell RNA sequencing, including size-factor normalization (Robinson and Oshlack) and variance-stabilizing transformations (Hafemeister and Satija), assume that technical effects can be estimated independently of the biological model of interest. Even normalization procedures developed specifically for spatial transcriptomics primarily target spatial variation in library size while treating adjusted values as fixed inputs to downstream analyses (Salim et al.). Such staged approaches are reasonable when technical effects can be estimated independently of the inferential target. In ST data, however, technical and biological spatial components are often only partially separable (Bhuva et al.). When similar spatial patterns may be attributed either to a latent technical component or to a biological component of interest, normalization is no longer merely a preprocessing step. It becomes part of the inferential problem. In this setting, two-stage normalization may cause bias in downstream statistical analysis such as differential expression analysis.

From a statistical standpoint, spatial transcriptomic data are naturally modeled using hierarchical generalized linear models with spatially structured random effects. Conditional autoregressive (CAR) and intrinsic CAR (ICAR) priors are widely used for areal data to represent local spatial dependence (Besag; Besag et al.), while Gaussian process models provide continuous-domain representations of spatial correlation (Rasmussen and Williams). Through the stochastic partial differential equation link, ICAR/GMRF priors can be viewed as sparse discrete approximations to smooth Gaussian processes (Lindgren et al.), which motivates the modeling choice adopted here. Inference in hierarchicalYes spatial models has been extensively studied, including scalable approaches based on integrated nested Laplace approximation (Rue et al.). Joint modeling of latent spatial effects and covariate effects is particularly important when covariates exhibit spatial patterning, as spatial confounding may otherwise distort fixed-effect estimation (Reich et al.; Hodges and Reich). These considerations suggest that normalization and downstream inference in spatial transcriptomics may be more coherently addressed within a unified hierarchical framework that explicitly represents spatial technical components, rather than being performed in separate stages.

Motivated by this perspective, we propose **GPSNorm** (Gaussian Process Spatial Normalization), a Bayesian hierarchical modeling framework that treats normalization as the joint estimation of structured nuisance and biological effects. Rather than adjusting counts in a preliminary step and treating normalized values as fixed, the framework represents expression through a negative binomial latent Gaussian model in which latent technical components and biological effects of interest are estimated simultaneously. Joint estimation reduces the risk that biologically meaningful variation is absorbed during normalization and ensures that uncertainty in the decomposition of technical and biological effects is propagated into downstream inference. The model is formulated within a Bayesian latent Gaussian framework and is estimated using integrated nested Laplace approximation (INLA; (Rue et al.)), which enables efficient approximate inference for complex latent Gaussian models. To further improve scalability for large spatial graphs, we investigate area-of-interest (AOI) clustering as a dimension-reduction strategy. While we focus on differential expression as an illustrative task, the framework can be extended to accommodate additional covariates and other structured effects, including spatially varying gene patterns, through appropriate specification of the linear predictor.

We evaluate the proposed framework through simulation studies anchored to empirical count distributions and through application to three spatial transcriptomics datasets: a 10x Visium human dorsolateral prefrontal cortex dataset (Maynard et al.; Pardo et al., 2022), a NanoString GeoMx dataset, and characterizing COVID-19 pneumonitis (Cross et al.) a NanoString GeoMx Spatial Organ Atlas kidney dataset (NanoString Technologies, 2026). Across these settings, we assess recovery of structured technical effects, estimation accuracy, uncertainty propagation, consistency with biological expectations, and computational scaling feasibility under varying levels of spatial confounding.

## 2 GPSNorm Modeling Framework

### 2.1 Data Structure and Notation

Let *g* = 1, …, *G* index genes and *i* = 1, …, *N* index spatial units. Depending on the platform, a spatial unit may correspond to a Visium spot, a GeoMx area of interest, or another spatially indexed region. For each gene *g* and spatial unit *i*, let *y*_*gi*_ denote the observed transcript abundance. Let *L*_*i*_ denote a known offset for spatial unit *i*, typically representing library size or sequencing depth. Covariates associated with spatial unit *i* are collected in a vector ***z***_*i*_ ∈ ℝ^*p*^, which may encode biological contrasts such as disease group, tissue compartment, or damage severity as well as additional technical factors.

Each gene has a control type indicator

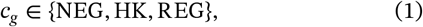

where NEG denotes negative control genes assumed not to exhibit biological variation, HK denotes housekeeping genes expected to show relatively stable expression across biological conditions, and REG denotes regular biologically variable genes. These annotations are used to help identify shared technical variation and to distinguish control features from genes used for biological inference.

Spatial dependence is represented through an undirected graph *G* = (*V, E*), where vertices *V* = {1, …, *N*} is the set of spatial units and edges *E* encodes neighborhood relationships among units within the same slide. Let *i* ∼ *j* denote adjacency, with adjacency matrix *W* = (*w*_*ij*_) and degree *d*_*i*_ = ∑_*j*_ *w*_*ij*_ . Spatial units are grouped by slide or batch; let *s*(*i*) ∈ {1, …, *S*} denote the slide associated with spatial unit *i*.

The goal of our normalization is to decompose expression into technical artifacts shared broadly across genes and gene-specific biological effects while preserving coherent uncertainty quantification for downstream inference. This decomposition rests on the identifying assumption that technical variation is shared across genes (captured by the slide and spatial terms), whereas gene-specific variation is biological. Gene-specific spatial biological structure is therefore not represented in the linear predictor and is treated as part of the residual.

### 2.2 Hierarchical Model Specification

Observed counts are modeled using a hierarchical generalized linear model in which technical and biological components are estimated jointly. For each gene *g* and spatial unit *i*,

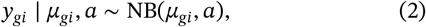

where *μ*_*gi*_ is the mean and *a* > 0 is the negative binomial size parameter, with variance

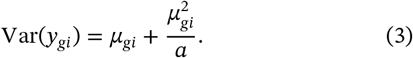

We adopt a log link,

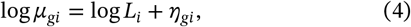

where *L*_*i*_ is treated as a known offset and *η*_*gi*_ is the linear predictor,

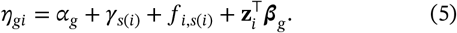

Here *α*_*g*_ is a gene-specific baseline effect, *γ*_*s*(*i*)_ is a slide-level effect, *f*_*i,s*(*i*)_ is a slide-specific spatial effect capturing structured technical variation within slide *s*(*i*), and ***β***_*g*_ are gene-specific regression coefficients. The offset log(*L*_*i*_) adjusts for global exposure, whereas *γ*_*s*(*i*)_ and *f*_*i,s*(*i*)_ capture residual slide-level and spatially structured variation not explained by the offset or measured covariates.

Control annotations are incorporated by omitting gene-specific biological parameters for negative controls For negative control genes we impose

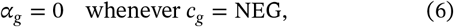

and do not estimate gene-specific covariate effects *β*_*g*_ for these genes, reflecting the assumption that negative controls do not carry gene-specific biological signal beyond shared technical components. This assumption follows standard control-based normalization approaches, where negative control or spike-in features are affected by technical factors but not by biological covariates, allowing them to anchor estimation of unwanted variation (Risso et al.; Evans et al.).

Under this constraint, the counts of negative controls beyond the known offset are explained by the shared slide-level and spatial components, so the negative controls anchor estimation of these technical terms and the dispersion parameter. This reduces confounding between technical and biological variation.

### 2.3 Prior Specification

#### 2.3.1 Independent Random Effects

We assign Gaussian priors to the independent components of the latent field. For genes not classified as negative controls, baseline effects, regression coefficients, and slide effects are assumed independent with

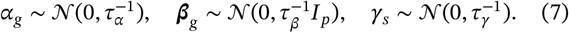

These priors provide regularization for gene-level baselines, biological covariate effects, and slide-level normalization effects.

#### 2.3.2 Spatially Structured Random Effects

Spatial dependence within each slide is modeled using an intrinsic conditional autoregressive (ICAR) prior. For slide *s*, let **f**_*s*_ = {*f*_*i,s*_ : *s*(*i*) = *s*}. Then

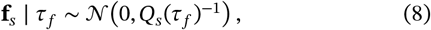

with precision matrix *Q*_*s*_(*τ*_*f*_) = *τ*_*f*_(*D*_*s*_ − *W*_*s*_), where *W*_*s*_ is the within-slide adjacency matrix and *D*_*s*_ is the corresponding diagonal degree matrix.

The full spatial precision matrix is block diagonal across slides, so that spatial effects are independent between slides, while a common precision parameter *τ*_*f*_ governs spatial smoothness across slides. A sum-to-zero constraint

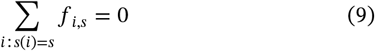

is imposed within each slide to identify the spatial field relative to the slide-level effect *γ*_*s*_ and gene-specific intercepts.

The ICAR prior can be viewed as a sparse Gaussian Markov random field (GMRF) representation of local spatial dependence Continuous Gaussian processes define spatial dependence through covariance functions over ℝ^2^, but lead to dense covariance matrices. In contrast, ICAR models specify local conditional dependence through the graph Laplacian (*D* − *W*), which serves as a finite-difference approximation to the continuous Laplace operator. Through the stochastic partial differential equation representation of Matérn Gaussian processes (Lindgren et al.), GMRFs arise as discrete representations of spatially smooth Gaussian processes. The resulting sparse precision structure enables scalable computation for large spatial graphs.

#### 2.3.3 Hyperpriors

By default, the R-INLA package (Lindgren and Rue) assigns diffuse Gamma hyperpriors to precision parameters. For a generic precision parameter *τ*, the prior is specified as

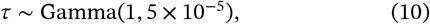

where the second parameter denotes the rate. This specification is used for the gene-specific baseline precision *τ*_*α*_, the gene-specific differential effect precision *τ*_*β*_, the slide-effect precision *τ*_*γ*_, and the spatial ICAR precision *τ*_*f*_ . Thus, the corresponding random-effect variances are primarily data driven.

Also following R-INLA recommendation, negative binomial size parameter *a* is assigned a penalized complexity marginal Gamma prior (Simpson et al.; Lindgren and Rue). This hyperprior serves to regularize the negative binomial likelihood towards a Poisson model (*a* → ∞). This way, additional overdispersion beyond Poisson variability is only introduced when supported by the data.

### 2.4 Latent Gaussian Representation

The proposed model belongs to the class of latent Gaussian models (LGMs) (Rue et al.). This follows directly from the linear predictor structure and the Gaussian specification of all latent components. Stack the observations *y*_*gi*_ into **y** and define the stacked linear predictor ***η*** = {*η*_*gi*_ }. Let the latent field be

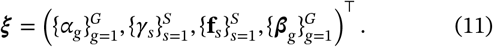

Because the predictor is additive in these components, there exists a design matrix *A* such that ***η*** = *A****ξ***, where *A* encodes gene indexing, slide membership, spatial-unit-to-slide mapping, covariates ***z***_*i*_, and the negative-control constraint on *α*_*g*_.

Conditional on the hyperparameters, the independent components, the independent components of ***ξ*** are Gaussian. Gene baselines *α*_*g*_, slide effects *γ*_*s*_, and regression coefficients ***β***_*g*_ are exchangeable Gaussian random effects with diagonal precision matrices determined by (*τ*_*α*_, *τ*_*γ*_, *τ*_*β*_ ), while each spatial field **f**_*s*_ follows an ICAR prior with precision matrix *τ*_*f*_(*D*_*s*_ − *W*_*s*_). Since independent Gaussian vectors remain jointly Gaussian, it follows that

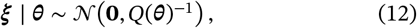

where ***θ*** = (*a, τ*_*α*_, *τ*_*β*_, *τ*_*γ*_, *τ*_*f*_) and *Q*(***θ***) is a block-diagonal precision matrix with blocks corresponding to gene baselines, slide effects, spatial fields, and regression coefficients. The spatial block itself is block diagonal across slides, with blocks *τ*_*f*_(*D*_*s*_ − *W*_*s*_), which are sparse because nonzero off-diagonal entries occur only for neighboring spatial units. Independence across hierarchical components and across slides therefore yields a sparse joint precision matrix.

Conditional on (***η***, *a*), observations are independent under the negative binomial likelihood, so the joint model factorizes as

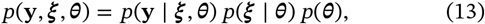

which matches the defining structure of a latent Gaussian model: a conditionally independent likelihood, a Gaussian latent field governed by a low-dimensional hyperparameter vector, and a sparse precision matrix (Rue and Held; Lindgren et al.).

### 2.5 Posterior Estimation

The latent Gaussian structure described above is computationally advantageous because posterior inference can be reduced to operations involving the sparse precision matrix of the latent field. Direct Markov chain Monte Carlo (MCMC) sampling in this setting would require repeated high-dimensional updates of the latent field, whose dimension grows with the number of genes and spatial units. Because ***ξ*** ∣ ***θ*** is Gaussian with sparse precision matrix *Q*(***θ***), deterministic approximations can exploit sparse matrix factorization to obtain accurate marginal posterior distributions.

Integrated nested Laplace approximation (INLA) (Rue et al.) provides a computational framework tailored to latent Gaussian models. INLA proceeds by first approximating the marginal posterior of the hyperparameters using nested Laplace approximations that integrate out the Gaussian latent field. Conditional on ***θ***, the posterior of ***ξ*** is approximated by a Gaussian distribution obtained via sparse Cholesky factorization of *Q*(***θ***) combined with a second-order Taylor expansion of the log-likelihood around its mode. Marginal posterior distributions of latent components are then obtained by numerical integration over ***θ***.

The computational efficiency of INLA in this setting arises from two structural features of the model. First, the hyperparameter vector ***θ*** = (*a, τ*_*α*_, *τ*_*β*_, *τ*_*γ*_, *τ*_*f*_) is low-dimensional, so numerical integration over ***θ*** is feasible. Second, the latent precision matrix is sparse due to the ICAR spatial structure and independence across hierarchical components. Sparse Cholesky factorization scales approximately linearly in the number of nonzero elements of *Q*(***θ***), making computation substantially more efficient than methods requiring dense covariance matrices, such as direct Gaussian process inference. In particular, replacing a dense spatial covariance matrix with an ICAR precision matrix avoids the cubic computational complexity in the number of spatial units that arises from inversion of dense Gaussian process covariance matrices. The cost of inference is instead governed by the sparsity pattern of the graph Laplacian, which depends on local neighborhood structure rather than total graph size. This makes latent Gaussian inference particularly well suited to spatial transcriptomic datasets where each spatial unit interacts only with a small number of neighbors.

### 2.6 Inference Procedures

Posterior inference is based on the marginal distributions returned by INLA. Posterior summaries such as the mean, median, standard deviation, and credible intervals are available for each component of the model, including gene-specific effects, spatial effects, slide effects, and dispersion parameters. Because normalization effects are included explicitly in the model, their posterior distributions can be examined individually rather than treated as fixed preprocessing adjustments. These quantities provide interpretable summaries of experimental variability, allowing researchers to assess batch effects, spatial artifacts, and sequencing depth differences across samples or tissue sections. Explicit modeling of technical effects is widely recognized as important in high-throughput transcriptomic experiments, where unaccounted sources of variation may confound biological interpretation (Lähnemann et al.; Haghverdi et al.).

Posterior inference may also be performed on quantities derived from the linear predictor. We consider the decomposition

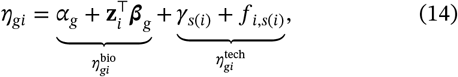

where the intercept, gene baseline, and regression effects are treated as biological, while slide and spatial terms are treated as technical. Here, 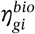 denotes the component retained for biological interpretation, whereas 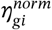 denotes the residual slide-level and spatial normalization component. This decomposition is not unique, and the choice of which terms are considered technical may depend on the scientific objective. Because all effects are estimated jointly, adjusted predictors can be constructed without refitting the model, allowing alternative normalization choices to be explored within the same fitted model.

Derived predictors can be projected back to the count scale to obtain normalized expression values. One approach computes an adjusted posterior-median count,

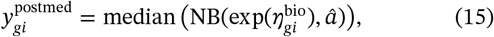

where *â* is the estimated dispersion parameter. This projection yields a smooth estimate of expression that reflects the expected biological signal after removing selected technical effects and is useful for visualization and model-based summaries.

An alternative approach constructs normalized counts using a percentile-adjusted count (PAC) transformation. Let *y*_*gi*_ denote the observed count and define

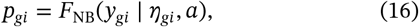

where *F*_NB_ is the negative binomial cumulative distribution function. The adjusted count is then obtained by

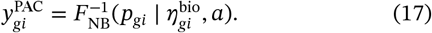

This transformation preserves the rank and dispersion of the observed counts while modifying the mean structure to remove selected technical effects. Percentile-based normalization has been shown to be effective for spatial transcriptomics, where technical variation may be confounded with spatial structure, and adjusted counts can improve downstream analyses such as clustering or spatial domain identification (Salim et al.).

The posterior-median and PAC projections provide complementary representations of normalized expression. The posterior-median projection is determined entirely by the fitted model and is therefore well suited for visualization and interpretation of model effects. In contrast, the PAC transformation preserves the variability of the observed counts while removing selected normalization terms, making it more appropriate for downstream analyses that rely on count-scale variation, such as clustering, dimension reduction, or spatially variable gene detection. Both representations are derived from the same fitted model and can be constructed without refitting, allowing the form of normalized expression to be chosen according to the goals of the analysis.

## 3 Propagation of Normalization Uncertainty

### 3.1 Theoretical Motivation

Differential expression (DE) analyses in spatial transcriptomics are commonly performed using a two-stage workflow in which normalization is estimated first and treated as fixed in a subsequent DE model. In this framework, normalization components, such as library-size adjustments, slide effects, or spatial correction terms, are estimated from the data and then incorporated as offsets or covariates without accounting for their estimation uncertainty. By contrast, GPSNorm estimates normalization and DE jointly within a single hierarchical model.

To formalize this, let *β* denote the DE parameters of interest and let *u* denote latent normalization components, such as spatial and slide effects. In a staged approach, normalization is treated as fixed, *û*, and inference proceeds as

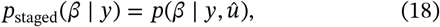

whereas the joint model integrates over the uncertainty in *u*,

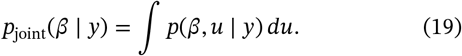

The consequences of this distinction follow from the law of total variance,

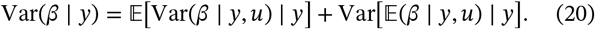

The first term represents uncertainty in the DE parameters conditional on a fixed normalization field, while the second term captures additional variability arising from uncertainty in the normalization components themselves. A staged analysis that conditions on a single estimate *û* effectively ignores the second term, because inference is performed as if the normalization were known.

The magnitude of this additional variance depends on the relationship between the DE covariates and the normalization structure. When group labels are approximately orthogonal to spatial and slide effects and the posterior uncertainty in *u* is small, the second term is negligible and staged and joint approaches yield similar results. However, when DE covariates are statistically associated with latent spatial or batch effects, multiple decompositions of the linear predictor may be consistent with the data, and the posterior mean of *β* can vary across plausible normalization fields. In this setting, conditioning on a single normalization estimate can lead to underestimation of uncertainty and anti-conservative inference. Joint estimation is therefore adaptive: it increases posterior uncertainty when normalization and DE are not fully separable, while reducing to conventional behavior when normalization uncertainty is negligible.

### 3.2 Empirical Demonstration Under Spatial Confounding

To illustrate the practical consequences of this distinction, we constructed a reduced spatial simulation that isolates the normalization–DE interaction while preserving the local adjacency structure. This adjacency structure was used within an ICAR prior consistent with the spatial component of the full model. Counts were generated according to

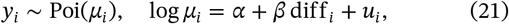

where dif f _*i*_ = 1(*x*_*i*_ > 0.5) defines a spatially structured group contrast and *u*_*i*_ = *σ*_*u*_ *ũ* _*i*_ is a smooth spatial field generated as a standardized cumulative sum of Gaussian increments. The spatial standard deviation *σ*_*u*_ controls the strength of latent structure; when *σ*_*u*_ = 0, the spatial field is exactly zero, providing a non-confounded baseline.

The true DE effect was set to *β* = 0 to enable direct assessment of interval calibration. Under this null configuration, nominal 95% credible intervals should contain the true value in approximately 95% of repeated datasets; deviations therefore reflect variance miscalibration rather than power limitations or estimation bias.

Figure 1 (left) displays the average 95% credible interval width for *β* as a function of spatial variance *σ*_*u*_. At *σ*_*u*_ = 0, joint and two-stage approaches coincide, consistent with the absence of normalization uncertainty. As spatial variance increases, credible intervals under joint estimation widen monotonically, reflecting the second term in Eq. 20. In contrast, interval widths under the two-stage approach remain nearly constant across spatial variance levels.

**FIGURE 1.**
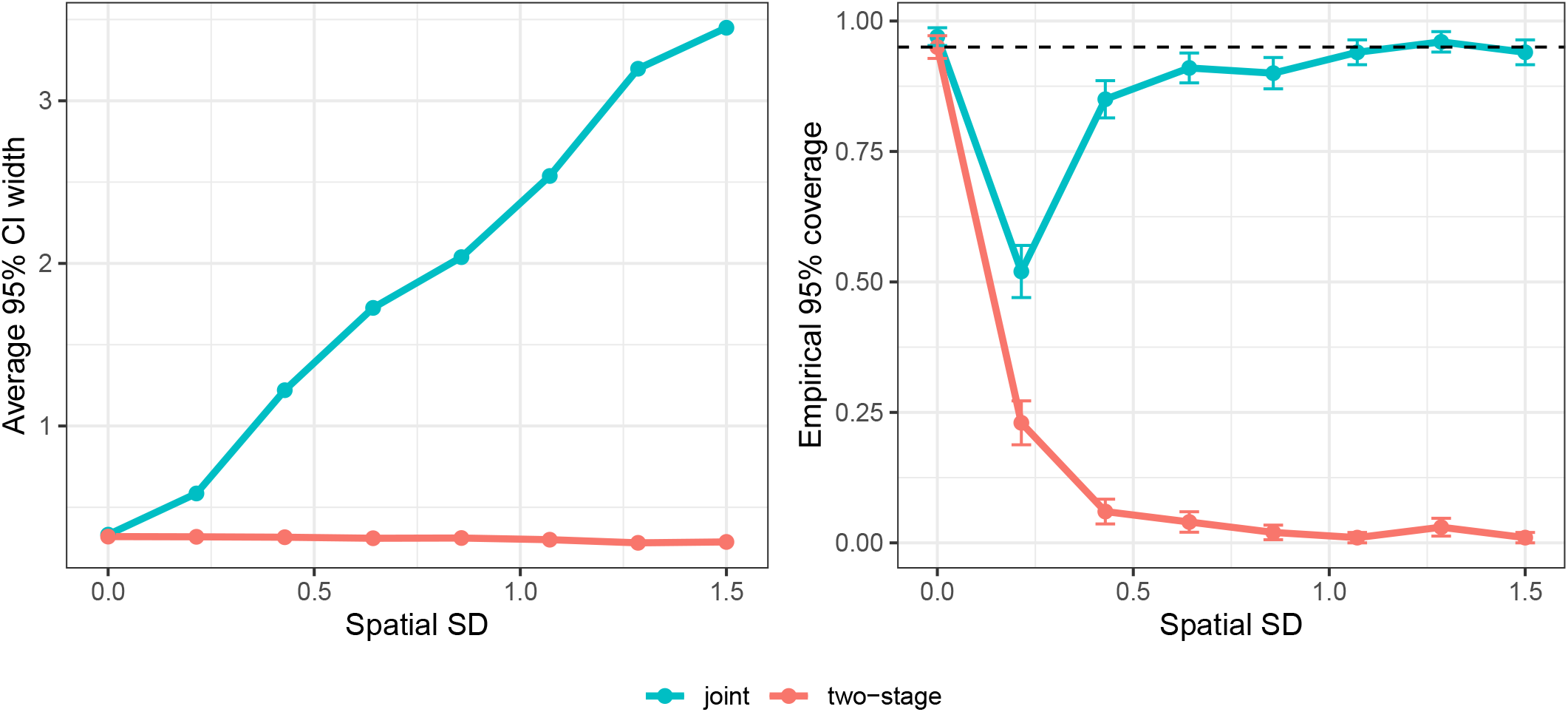
Interval width and coverage under two-stage normalization versus Bayesian joint estimation, in the reduced spatial simulation (Eq. 21) with true *β* = 0. Left: average 95% credible-interval width for *β* as a function of the spatial standard deviation *σ*_*u*_ . Right: empirical 95% coverage; the dashed line marks the nominal 0.95 level. Points are means over 100 replicates per *σ*_*u*_ level; error bars denote ±1 SE based on a binomial distribution.

Figure 1 (right) reports empirical 95% coverage across 100 simulation replicates per variance level. Joint estimation maintains coverage near the nominal level for across most levels of *σ*_*u*_, with a transient drop at low spatial variance, whereas coverage under two-stage normalization deteriorates as spatial variance increases, approaching zero under strong confounding. These results demonstrate that failure to propagate normalization uncertainty produces systematically underestimated DE variance and anti-conservative inference when DE covariates correlate with latent spatial structure.

Although this simulation isolates the spatial component, the variance decomposition in Eq. 20 applies to all latent normalization layers in the full hierarchical model, including slide effects and other structured components. Whenever DE covariates are not fully separable from these latent sources of variability, joint Bayesian estimation appropriately inflates uncertainty and preserves calibration, whereas conventional two-stage normalization cannot.

## 4 Simulation Study

### 4.1 Simulation Design

To evaluate performance under realistic yet controlled conditions, we constructed a generative framework that preserves empirical count characteristics while allowing systematic manipulation of technical and biological structure. The simulation design follows principles recommended in recent spatial transcriptomics benchmarking studies, which emphasize empirical calibration, structured confounding, and explicit spatial modeling to ensure meaningful method evaluation (Liang et al.).

#### 4.1.1 Empirical Calibration of Baseline Expression

Baseline expression and dispersion parameters are anchored to real data to preserve sparsity and heterogeneity. For each gene *g*, we estimate a baseline rate

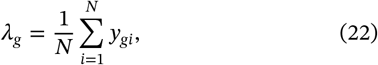

and simulate counts under a negative binomial model with gene-specific dispersion *θ*_*g*_,

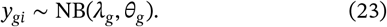

The negative binomial likelihood is widely used for RNA-seq data due to its ability to capture overdispersion beyond Poisson variability (Love et al.). Empirical anchoring ensures that simulated data reflect realistic variability in gene abundance and dispersion.

#### 4.1.2 Technical Variation

Library size variation is a dominant technical artifact in sequencing-based assays (Love et al.). Library size factors *L*_*i*_ are sampled from the empirical distribution observed in real datasets,

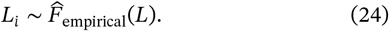

Slide-level technical effects are introduced to mimic batch variation shared across areas of interest (AOIs),

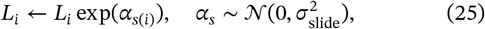

where *s*(*i*) indexes slides. Slide-level multiplicative shifts are a common source of variability in spatial transcriptomics data (Millard et al.).

Spatially structured technical effects are generated using a smooth radial basis function Gaussian process, 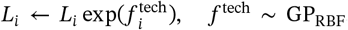, which induces spatial bias unrelated to biology (Qian et al.; Lindgren et al.).

#### 4.1.3 Biological Signal Components

Differential expression (DE) effects are introduced through group-specific log fold changes,

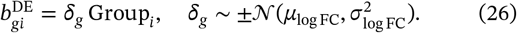

To reflect structured biological heterogeneity, domain-specific effects are added,

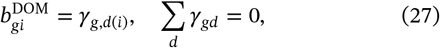

where *d*(*i*) denotes anatomical domains.

Spatially variable genes are simulated using gene-specific spatial fields,

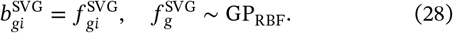

Overlap between signal categories is controlled by

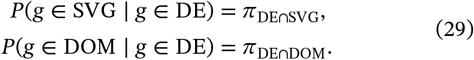

Varying these probabilities generates regimes of increasing biological confounding (Liang et al.).

#### 4.1.4 Compositional Structure and Background Noise

To reflect the compositional nature of sequencing data, gene-level proportions are constructed as

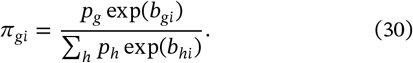

Background contamination is introduced through

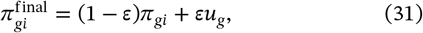

where *u*_*g*_ is a gene-level noise distribution and *ε* controls contamination intensity.

Counts are then generated as

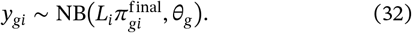

This hierarchical simulation framework allows independent control of slide-level, spatial, and biological structure while preserving empirical count characteristics, enabling rigorous evaluation of normalization and downstream inference under varying degrees of technical and biological confounding.

### 4.2 Simulation Scenarios

All simulations were generated using the hierarchical framework described in Section 4.1. Across scenarios, baseline expression, dispersion, compositional structure, and background contamination were held fixed. Scenarios differ only in the degree of biological overlap and in the magnitude of technical variation. This structured grid allows us to isolate identifiability challenges arising from biological confounding, technical confounding, and their interaction. The defining parameter settings for each scenario are summarized in Table 1.

**TABLE 1.**
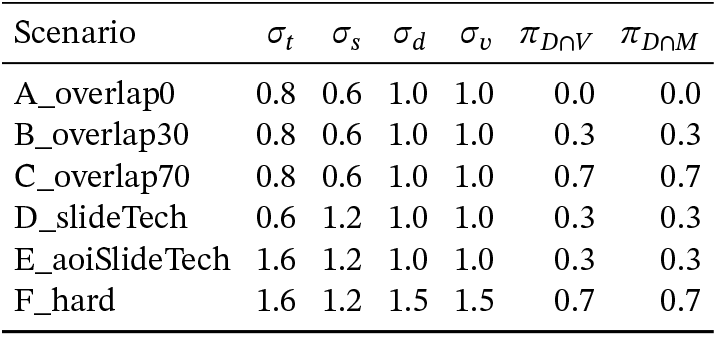
Simulation scenario parameters. *σ*_*t*_ = amplitude of the technical spatial GP. *σ*_*s*_ = slide-effect SD. *σ*_*d*_ = magnitude of the spatial domain effect. *σ*_*v*_ = amplitude of the SVG GP. *π*_*D*∩*V*_ = probability that a gene is DE and SVG. *π*_*D*∩*M*_ = probability that a DE gene has a domain effect. All other parameters held fixed across scenarios.

We first vary the degree of overlap between differential expression (DE), domain-specific expression (DOM), and spatially variable genes (SVGs) while maintaining moderate technical effects.

- **A_overlap0** represents a clean baseline with no overlap among DE, DOM, and SVG genes. This regime provides an upper bound on achievable recovery when biological signals are separable.
- **B_overlap30** introduces moderate biological entanglement, with 30% overlap between DE genes and structured signals (DOM and SVG). This setting reflects realistic scenarios in which spatial and domain effects partially coincide with group contrasts.
- **C_overlap70** imposes strong biological confounding, with 70% overlap. In this regime, genes exhibiting differential expression frequently also show domain-specific or spatial structure, substantially increasing identifiability difficulty.

Increasing biological overlap primarily stresses downstream differential expression estimation, as structured biological signals may be partially absorbed by normalization procedures or confounded with technical components. To isolate the impact of technical variation, we next vary the magnitude of slide-level and spatial technical effects while holding biological overlap at a moderate level.

- **D_slideTech** increases the variance of slide-level effects *σ*_slide_, producing strong batch-like shifts shared across AOIs within a slide.
- **E_aoiSlideTech** increases both slide-level and spatial technical variance, introducing smooth AOI-level technical structure in addition to batch effects.

These regimes primarily test the ability of normalization methods to recover structured technical spatial fields without distorting biological effects.

- **F_hard** combines strong biological overlap (70%) with strong slide-level and spatial technical variation. The difficultly in compounded by additionally increase the size of the un-modeled domain and SVG effects. This scenario represents a maximal confounding regime in which technical and biological structure are both pronounced and partially aligned.

This scenario grid enables systematic evaluation of method performance across progressively challenging regimes, disentangling the effects of biological overlap and technical confounding on spatial normalization and downstream inference.

### 4.3 Simulation Results

We evaluated performance under the simulation regimes described above using two complementary criteria. First, we examined recovery of the simulated technical spatial field to determine whether the normalization model accurately captures structured technical variation. Because the true technical field is known in simulation, recovery can be assessed by comparing the estimated spatial effect to the generating field. Second, we evaluated downstream differential expression (DE) performance to determine whether biological signal is preserved after normalization.

#### 4.3.1 Recovery of Technical Spatial Structure

The normalization model includes a spatial random effect intended to capture structured technical variation across spatial units. Because the simulation framework explicitly generates a technical spatial field, recovery of this component can be evaluated directly by comparing the estimated spatial effect to the true generating field.

Figure 2 illustrates this behavior for a representative simulation replicate under scernario /*F*_*hard*_. The left panel shows the true technical spatial field used during data generation, while the right panel shows the spatial effect estimated by the normalization model. The estimated field closely reproduces the large-scale spatial structure of the generating process, indicating that the model successfully identifies structured technical variation across spatial units.

**FIGURE 2.**
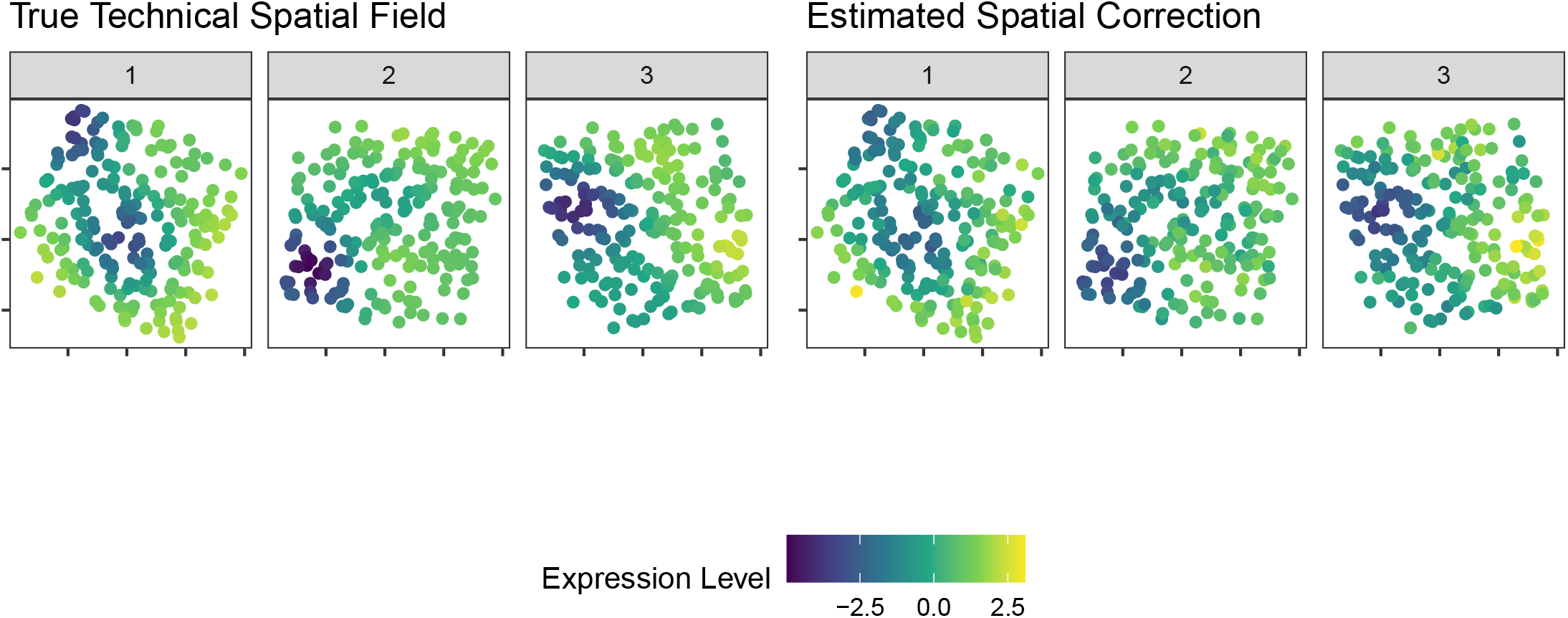
Example recovery of the simulated technical spatial field from a single simulation replicate under scenario F_hard. Left: true field *f*^tech^ used in data generation; right: GPSNorm-estimated spatial effect 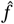 across the three simulated slides. Colors display the attributable spatial effect on expression level on a shared log scale.

While Figure 2 provides a visual illustration, we also examined quantitative recovery under the most challenging simulation regime (F_hard), which combines strong biological overlap with strong slide-level and spatial technical effects. For each replicate, recovery was measured as the correlation between the estimated spatial effect and the true technical spatial field. Across 25 independent simulation runs, correlations remained consistently high (0.921 ± 0.021), indicating stable recovery of the technical spatial structure even under substantial biological and technical confounding.

Together, these results suggest that the spatial normalization component accurately captures structured technical artifacts while preserving the underlying spatial organization of the technical field. Having established that the model can recover technical spatial structure, we next examine the impact of normalization on downstream differential expression estimation.

#### 4.3.2 Differential Expression Performance

We next evaluated downstream differential expression (DE) performance after normalization. Because the simulation framework generates known log fold-changes, estimation accuracy can be assessed directly by comparing estimated effects to their true values. We consider three criteria: bias of estimated log fold-changes, root mean squared error (RMSE), and ranking performance.

Figure 3 summarizes bias and RMSE of estimated log fold-changes across simulation scenarios. Each point represents the mean bias and RMSE across simulation replicates for a normalization method and scenario. GPSNorm exhibits minimal bias and reduced RMSE relative to competing approaches. The improvement becomes more pronounced as technical complexity increases, particularly in scenarios with strong slide-level or spatial technical variation. These gains arise from the hierarchical Bayesian formulation, which shares information across genes and spatial units when estimating technical effects and gene-level parameters.

**FIGURE 3.**
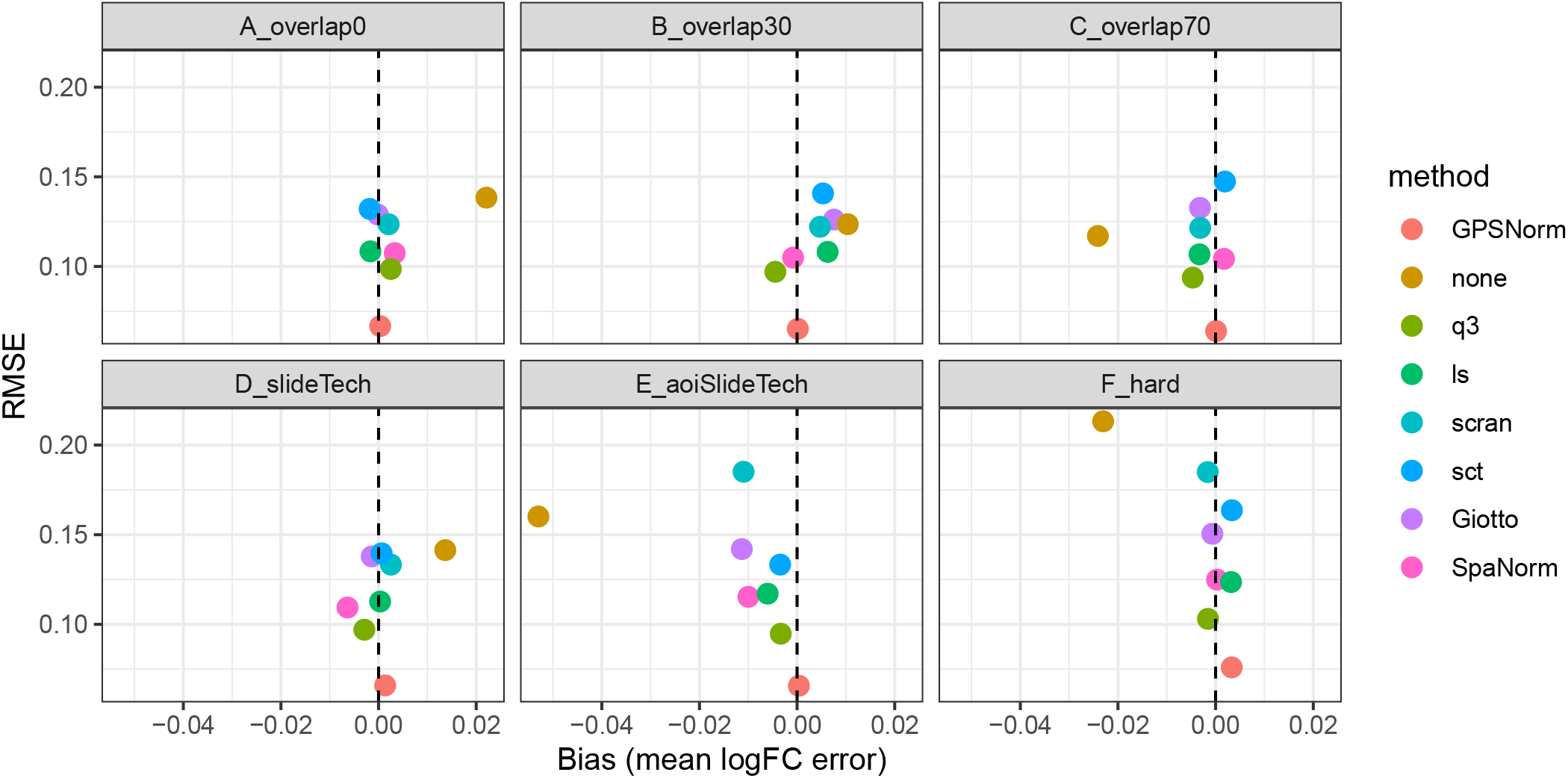
Bias and RMSE of differential expression effect size estimates. Mean log fold-change bias (x-axis) and RMSE (y-axis) across 30 simulation replicates for six simulation scenarios. Each point represents the mean performance of a normalization method. The dashed vertical line indicates zero bias. GPSNorm exhibits minimal bias and reduced RMSE across all settings.

Figure 4 provides a complementary summary of estimation error across simulation replicates. Bars show the mean RMSE of log fold-change estimates for each method and scenario, with error bars representing standard error across simulation runs. GPSNorm achieves the lowest RMSE in all scenarios and also exhibits reduced variability across replicates, reflecting the stabilizing effect of hierarchical information sharing.

**FIGURE 4.**
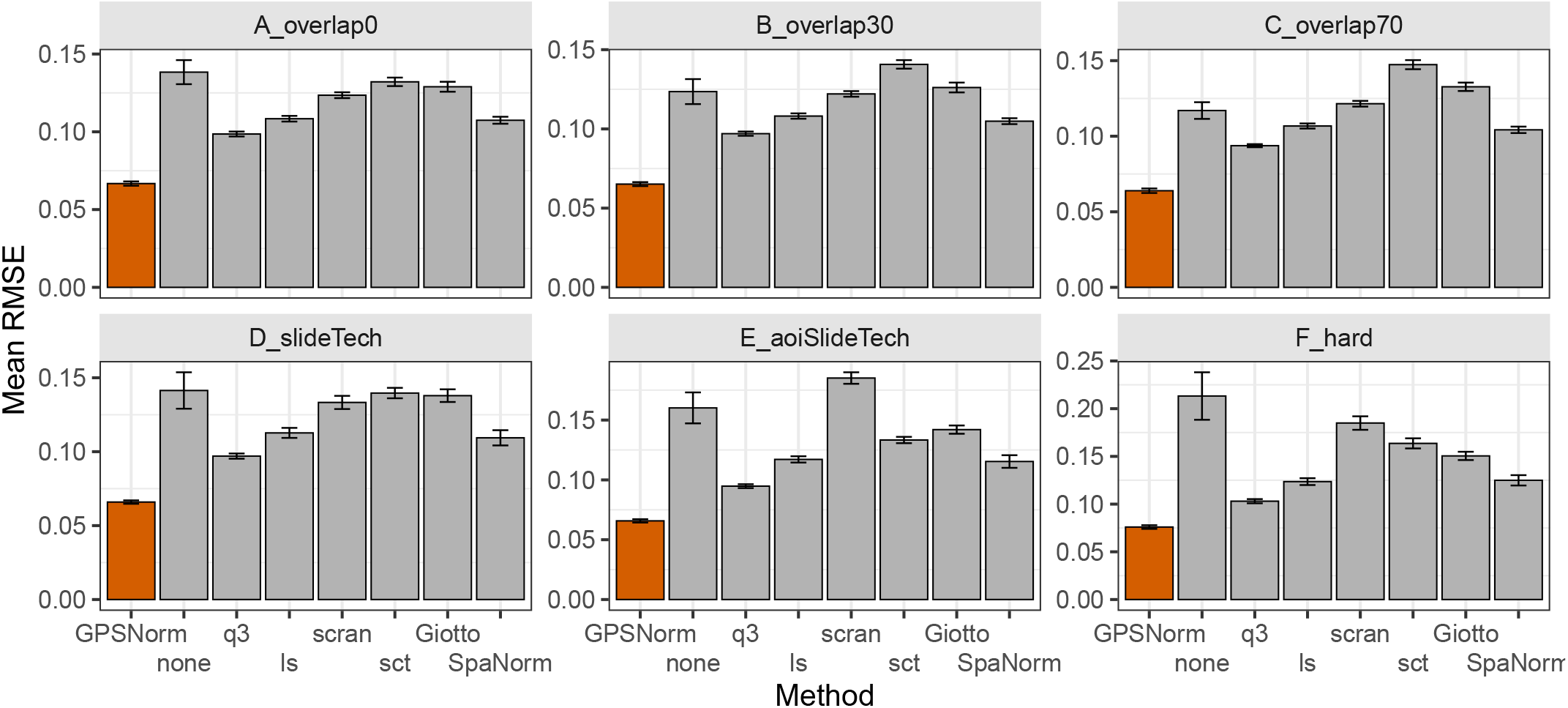
Mean RMSE across simulation replicates. Bars show the mean RMSE of log fold-change estimates across 30 simulation replicates for each normalization method and scenario. Error bars denote standard error across replicates. The lowest bar under each scenario is highlighted in orange. GPSNorm achieves the lowest RMSE in all scenarios, with larger improvements under increasing technical complexity.

Finally, Figure 5 evaluates the ability of each method to correctly rank DE genes using the area under the ROC curve (AUC). Ranking performance is similar across methods in simpler regimes with limited confounding. As technical variation increases, GPSNorm maintains higher AUC relative to alternative normalization approaches. This behavior reflects the regularizing effect of the model priors, where shrinkage toward the null stabilizes estimates for genes with little or no true effect and reduces noise-driven fluctuations.

**FIGURE 5.**
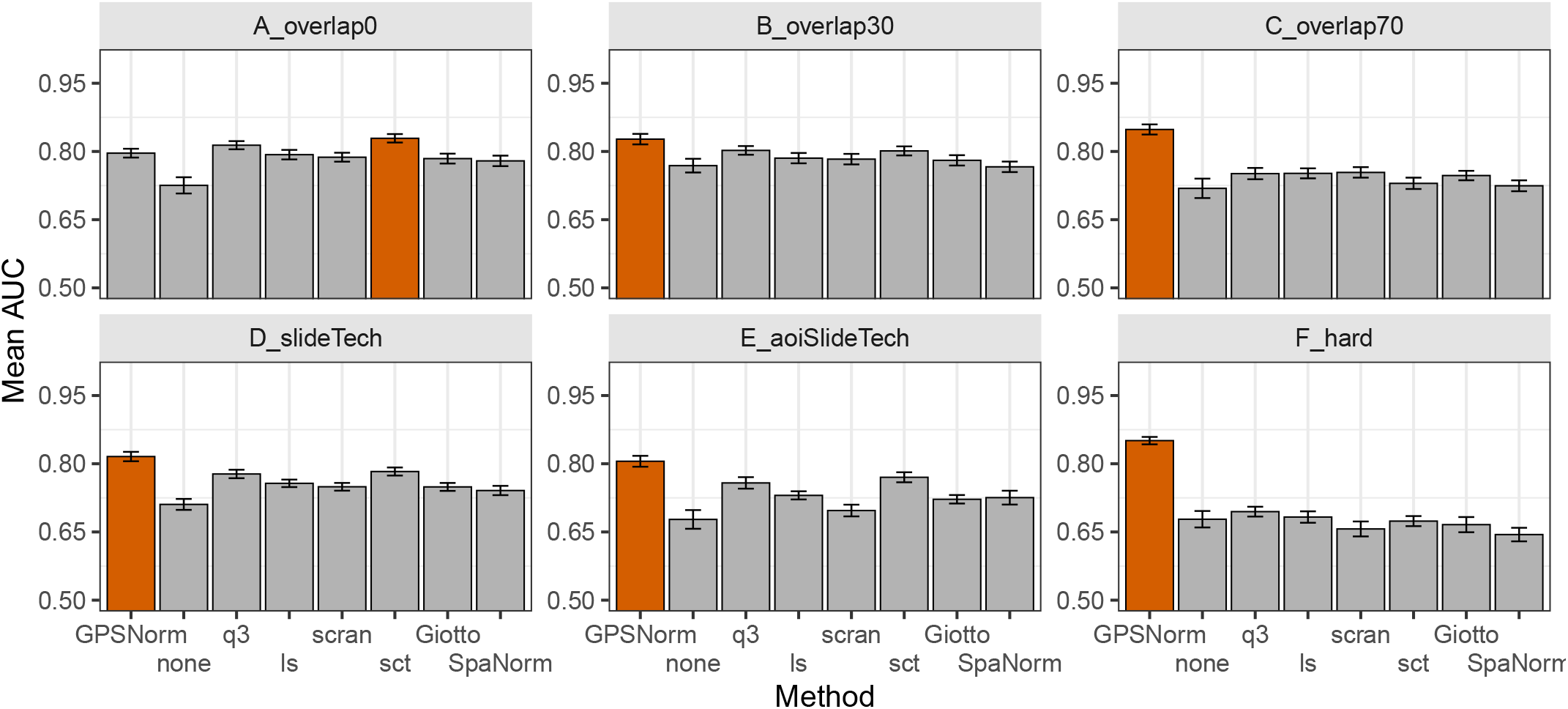
Ranking performance for differential expression detection across simulation scenarios. Bars show mean AUC across 30 simulation replicates for each normalization method. Higher values indicate better agreement with the true ordering of differential expression effects. The highest bar under each scenario is highlighted in orange. GPSNorm maintains competitive performance in simple settings and improves ranking under technical confounding.

Overall, these results demonstrate that joint Bayesian modeling improves the accuracy and stability of differential expression estimates. By borrowing information across genes and spatial units, the model more effectively separates technical spatial variation from biological signal, leading to reduced estimation error and improved identification of truly differentially expressed genes.

## 5 AOI Clustering and Computational Scaling

For reduced-resolution analyses, AOIs were aggregated only for model fitting; normalized values and downstream DE estimates were then projected back to the original AOI resolution. Aggregation was performed independently within each slide to avoid merging AOIs across slides. AOIs were visited in a seed-controlled random order, and each unassigned AOI initiated a cluster that was expanded using unassigned nearest neighbors until the target size was reached or no eligible neighbors remained.

After clustering, counts and library sizes were summed within each gene-by-cluster stratum, cluster coordinates were defined as the mean standardized coordinates of member AOIs, and binary group labels were assigned by majority rule. Gene class indicators were propagated to the cluster level, and each slide-cluster pair was assigned a unique AOI identifier for model fitting. Explicit checks were used to ensure complete AOI-to-cluster mappings, finite offsets, unique clustered AOIs, and unique gene-by-cluster observations.

We assessed the computational and statistical effects of AOI aggregation in both simulation (Table 2) and the Spatial Organ Atlas kidney dataset (Table 3). In simulation, moderate aggregation reduced runtime by roughly 50%, and more aggressive aggregation yielded approximately a four-fold speedup. In the kidney dataset, runtime decreased by more than five-fold at the coarsest resolution. These reductions are consistent with the dependence of inference cost on spatial graph dimension and indicate that aggregation provides a practical approach for scaling the model.

**TABLE 2.** Computational scaling and differential expression (DE) consistency under AOI aggregation in simulation. Results are shown for Scenario F_hard across 25 replicates. Rows correspond to the full-resolution fit and to 0.5x and 0.25x aggregation levels. AOI counts and runtime in seconds are averaged across replicates. RMSE and Spearman correlations are computed relative to the full-resolution model, so reference values are reported as NA. Top 10% denotes the Spearman correlation among the 10% of genes with the largest absolute logFC estimates under the full-resolution model.

| Res. | AOIs | Runtime (s) | RMSE | Spearman | Top 10% |
| --- | --- | --- | --- | --- | --- |
| 1.0x | 1206.00 | 236.78 | NA | NA | NA |
| 0.5x | 528.96 | 127.04 | 0.0548 | 0.5846 | 0.8227 |
| 0.25x | 347.16 | 62.90 | 0.0660 | 0.4627 | 0.7320 |

**TABLE 3.**
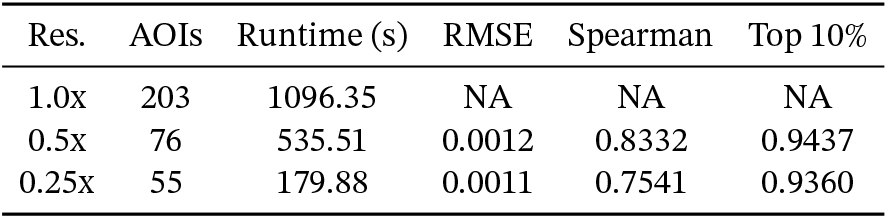
Differential expression (DE) consistency on the Spatial Organ Atlas kidney dataset across AOI aggregation levels. Rows correspond to the full-resolution fit and to 0.5x and 0.25x aggregation levels. For a single dataset, AOI aggregation is seeded to be deterministic, so the AOI counts are exact. Runtime is in seconds. RMSE and Spearman correlations are computed relative to the full-resolution model, so reference values are reported as NA. Top 10% denotes the Spearman correlation among the 10% of genes with the largest absolute logFC estimates under the full-resolution model.

To evaluate stability, we compared log fold-change (logFC) estimates to the full-resolution fit. In simulation, magnitude differences remained small (on the order of a few hundredths), suggesting limited distortion of effect sizes. Although global rank correlations decreased under stronger aggregation, agreement among the top-ranked genes remained substantially higher, indicating that most variability occurred among near-null effects.

In the kidney dataset, magnitude differences were negligible (on the order of 10^−3^), and rank correlations remained moderate to strong, with high agreement among the top 10% of genes. Because the full-resolution model serves as a reference rather than ground truth, these comparisons reflect consistency rather than accuracy.

Overall, AOI aggregation provides substantial computational gains while largely preserving DE magnitudes and the ordering of high-signal genes.

## 6 Real Data Applications

We next evaluate the performance of GPSNorm on several real spatial transcriptomics datasets spanning two widely used platforms: 10x Genomics Visium and NanoString GeoMx DSP. These datasets represent complementary experimental settings that highlight different challenges encountered in spatial transcriptomics analysis. In particular, they illustrate the ability of spatially aware normalization to recover known spatial biological structure, stabilize differential expression inference in small spatial studies, and preserve biologically validated marker programs across heterogeneous tissue compartments.

Unlike simulation studies where the ground truth is known, real data applications must be assessed through biological plausibility and consistency with established spatial expression patterns. Accordingly, the analyses presented here focus on three practical evaluation criteria. First, we examine whether normalization preserves spatial expression patterns for well-characterized marker genes. Second, we evaluate the stability of downstream differential expression analysis in settings with limited numbers of spatial observations. Third, we assess whether normalization preserves expected transcriptional contrasts between well-defined anatomical compartments.

The following datasets illustrate these behaviors in practice. We first examine a human dorsolateral prefrontal cortex (DLPFC) Visium dataset to assess recovery of spatially structured gene expression. We then analyze a GeoMx COVID-19 lung damage dataset to evaluate differential expression performance in a small-sample spatial study. Finally, we examine the GeoMx Spatial Organ Atlas kidney dataset to evaluate preservation of compartment-specific transcriptional programs using curated marker gene sets.

### 6.1 10x Visium Human DLPFC Dataset

We first evaluate normalization behavior using the 10x Genomics Visium human dorsolateral prefrontal cortex (DLPFC) dataset (Maynard et al.; Pardo et al., 2022). The DLPFC exhibits well-characterized laminar organization and a clearly defined white matter (WM) region, making it a useful benchmark for assessing whether normalization preserves biologically meaningful spatial structure. In particular, oligodendrocyte-associated genes are expected to exhibit strong enrichment within the WM relative to surrounding cortical layers.

To illustrate this behavior, we examine the spatial expression pattern of **MOBP** (myelin-associated oligodendrocyte basic protein), a canonical oligodendrocyte marker gene. Raw spatial transcriptomics counts often exhibit substantial variation in sequencing depth across spots. As a result, apparent spatial patterns in the raw counts can be dominated by library size variation rather than biological signal (Salim et al.). In the unnormalized data, this technical variability obscures the expected WM enrichment of MOBP.

Normalization methods that do not incorporate spatial coordinates can recover signal enrichment near the boundary between white matter and cortex, but often fail to recover strong enrichment within the WM region itself (Salim et al.). In contrast, by explicitly modeling spatial variation, GPSNorm and SpaNorm detect clear enrichment of MOBP within the WM region as well as the expression gradient at the boundary.

Although absolute expression scales differ between methods due to model-based rescaling, the biologically relevant feature is the relative spatial contrast between tissue compartments, which is preserved. GPSNorm additionally provides posterior uncertainty for the spatial expression surface, allowing enrichment patterns to be interpreted probabilistically rather than relying on a single normalized estimate. Because normalization and downstream modeling are jointly estimated, uncertainty from normalization propagates through the posterior distribution.

Figure 6 (A) illustrates this behavior using the oligodendrocyte marker gene **MOBP** in the human DLPFC dataset. Unnormalized counts obscure the expected white matter (WM) enrichment due to strong library size heterogeneity. Deterministic normalization approaches partially recover the spatial pattern but treat normalized values as fixed. GPSNorm instead estimates a posterior spatial surface, shown through percentile-adjusted counts (PAC) and posterior median expression. Although GPSNorm may shift overall expression levels due to model-based scaling, biological interpretation depends on relative spatial contrast. The WM region remains enriched relative to cortical layers, demonstrating preservation of biologically meaningful signal.

**FIGURE 6.**
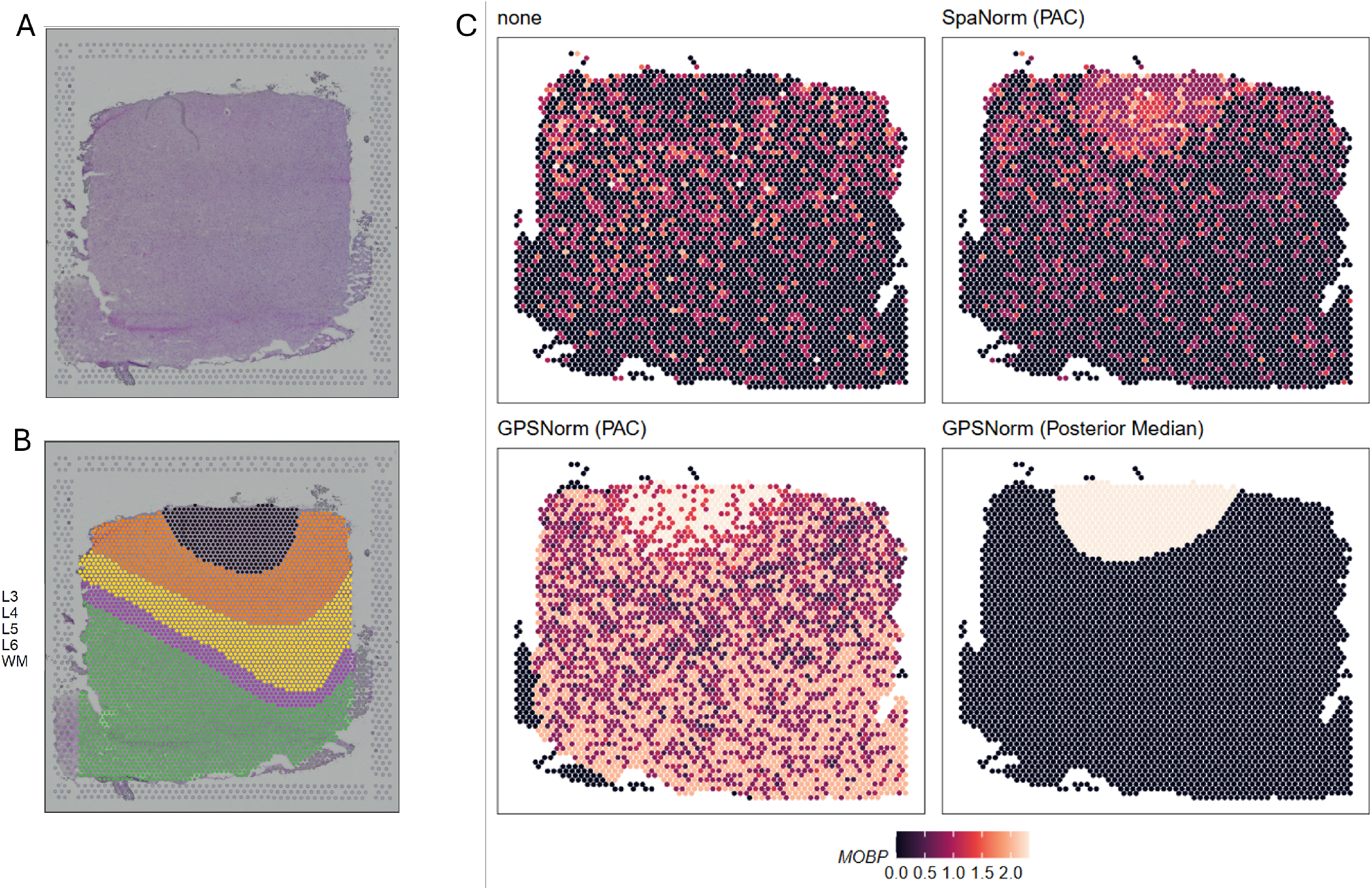
(A) Raw H&E pathology slide from the human DLPFC dataset (sample 151672). (B) Expert layer annotations (L3-L6, WM). (C) Spatial expression of the oligodendrocyte marker MOBP under four settings: unnormalized (none), SpaNorm (PAC), GPSNorm (PAC), and GPSNorm (posterior median). Unnormalized counts obscure white matter (WM) enrichment due to technical variation. Both SpaNorm and GPSNorm recover the signal within and at the boundary of the WM region. Color encodes relative MOBP expression on a shared log scale. GPSNorm preserves higher relative expression in WM compared to surrounding cortical layers.

### 6.2 GeoMx COVID-19 Lung Damage Dataset

We next evaluate normalization performance in a setting with a limited number of spatial observations using the NanoString GeoMx COVID-19 lung damage dataset (Cross et al.). This dataset contains 46 AOIs sampled from lung tissue sections exhibiting varying levels of pathological damage following SARS-CoV-2 infection. Each AOI was annotated according to the severity of tissue injury, enabling differential expression analysis comparing regions of **severe damage** against regions of **mild to moderate damage**. The dataset spans multiple slides and includes experimentally defined housekeeping (HK) genes and negative control probes, both of which are explicitly modeled by GPSNorm.

Differential expression analysis was performed by comparing AOIs labeled as severe damage to those labeled as mild or moderate damage. Each normalization method was applied directly to the expression data released by the original study through the Gene Expression Omnibus. No dataset-specific preprocessing or hyperparameter tuning was performed so that differences reflect normalization behavior rather than dataset-specific optimization. SpaNorm and scran were excluded because neither method produced valid estimates given the small number of AOIs.

Figure 7 summarizes the resulting differential expression analyses using volcano plots for each normalization method. Under GPSNorm, several chemokine genes known to be associated with severe COVID-19 lung inflammation are identified as strongly up-regulated in regions of severe damage, including **CCL18, CXCL10**, and **CCL2**, consistent with previous reports (Cross et al.). GPSNorm also identifies **SCGB1A1** as up-regulated in AOIs with milder damage, consistent with its known anti-inflammatory role in airway epithelial cells.

**FIGURE 7.**
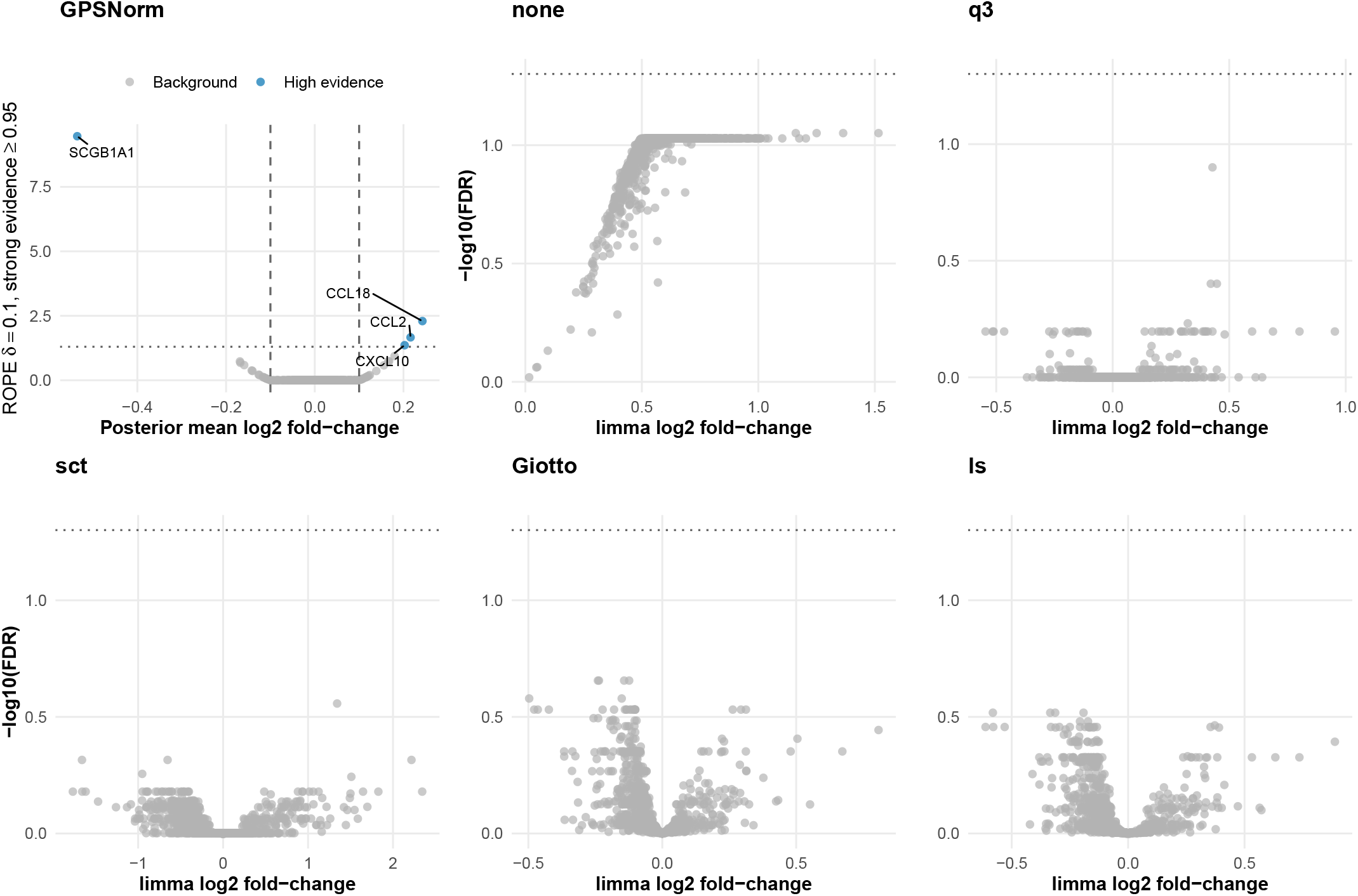
Differential expression between areas of severe and mild-to-moderate damage across normalization methods. Methods were applied directly to the NanoString GeoMx DSP dataset without additional dataset-specific preprocessing, quality-control adjustments, or hyperparameter tuning. GPSNorm differential expression is based on posterior inference using a region of practical equivalence (ROPE). Effects were deemed significant when *P*(|*β*_*g*_| > 0.10 ∣ *y*) ≥ 0.95. This corresponds to the threshold − log_10_(1 − 0.95) = 1.30 (marked by the horizontal dotted lines), analogous to the conventional − log_10_(0.05) cutoff used in FDR-based volcano plots. The GPSNorm ROPE-evidence axis and the limma − log_10_(FDR) axis are comparable as monotone evidence scales, with larger values indicating stronger evidence against a practically null effect, although they represent different inferential quantities. Only GPSNorm identified differentially expressed genes, including chemokines CCL18, CXCL10, and CCL2 in severely damaged regions, and SCGB1A1 in milder regions. SpaNorm and scran failed to produce valid estimates due to the small sample size.

In contrast, the remaining normalization approaches, each followed by a conventional limma test with FDR control, fail to identify statistically significant genes. This reflects the difficulty of differential expression analysis in spatial datasets with limited numbers of observations. When normalized expression values are treated as fixed quantities, uncertainty in normalization is not propagated into downstream inference, which can lead to unstable estimates or overly conservative results.

GPSNorm jointly models normalization and differential expression within a Bayesian framework, allowing information to be shared across genes, AOIs, and slides. The model also incorporates experimentally defined negative controls and housekeeping genes, which help anchor background expression and technical variation. These components stabilize differential expression estimates and allow biologically meaningful signals to be detected even when the number of AOIs is small.

These results suggest that Bayesian information sharing and explicit modeling of platform-specific control probes can substantially improve inference in small spatial transcriptomics studies. While this dataset demonstrates stability under small-sample conditions, it does not assess whether normalization preserves biologically expected transcriptional programs across tissue compartments. We therefore next examine a dataset with well-characterized marker genes to evaluate preservation of compartment-specific expression patterns.

### 6.3 GeoMx Spatial Organ Atlas Kidney Dataset

To assess differential expression (DE) performance on the GeoMx Spatial Organ Atlas kidney dataset, we curated marker gene sets representing the glomerulus and proximal tubule compartments. AOIs annotated as Cortical glomerulus or Juxtamedullary glomerulus were grouped as the positive class, and all other nephron segments were treated as the comparison group.

The glomerular marker set was derived from canonical podocyte and glomerular markers reported in renal biology studies, including **NPHS1, PODXL, SYNPO**, and **WT1** (Liu et al., b). These genes represent slit diaphragm structure, cytoskeletal specialization, and podocyte transcriptional identity. The proximal tubule marker set represents core solute transport and endocytic machinery characteristic of proximal tubular epithelial cells, including **LRP2, CUBN, AQP1, SLC22A6**, and **SLC22A8** (Bondue et al.).

Two complementary metrics were used. Mean evidence percentile measures how highly marker genes rank relative to the genome-wide DE distribution, providing a method-agnostic measure of statistical prioritization. Relative effect size enrichment quantifies the magnitude and direction of log fold-change for marker genes relative to the background distribution, assessing preservation of biological contrast.

Figure 8 summarizes the ability of each normalization method to prioritize known glomerular and proximal tubule marker genes. Glomerular markers rank highly across most methods, indicating that the dominant glomerular signal is robust to normalization choice, whereas larger differences appear for proximal tubule markers. GPSNorm maintains high ranking for both marker sets simultaneously, while several alternative normalization methods show reduced prioritization of tubule markers. Notably, the spatially aware SpaNorm does not show this advantage, suggesting that the gain reflects GPSNorm’s joint Bayesian modeling and control-gene anchoring rather than spatial awareness alone.

**FIGURE 8.**
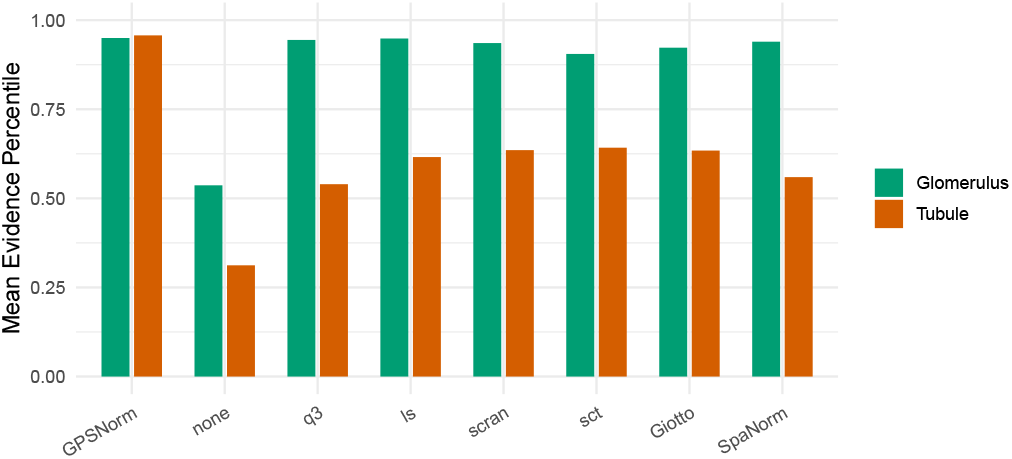
Marker-based evaluation of differential expression across normalization methods. Mean evidence percentile of curated glomerular and proximal tubule marker genes, reflecting their relative ranking within each method’s genome-wide DE results. Higher values indicate stronger prioritization of biologically expected markers.

Taken together, these real data analyses illustrate the practical advantages of spatially aware normalization and joint modeling in spatial transcriptomics studies. In the DLPFC Visium dataset, GPSNorm recovers known spatial expression patterns associated with white matter organization despite substantial library size heterogeneity. In the GeoMx COVID-19 dataset, the model stabilizes differential expression inference in a small spatial cohort, identifying biologically relevant chemokines and epithelial response genes not detected under alternative normalization approaches. In the Spatial Organ Atlas kidney dataset, GPSNorm preserves expected transcriptional contrasts between glomerular and tubular compartments while maintaining strong prioritization of curated marker genes.

Together, these results demonstrate that explicitly modeling spatial technical effects, slide structure, and platform-specific control probes improves both statistical robustness and biological interpretability across diverse spatial transcriptomics platforms.

## 7 Discussion

Spatial transcriptomics technologies have provided new opportunities to study gene expression in its native tissue context, but they also create a gap between the need to remove substantial technical variation, such as library size heterogeneity, slide-level batch effects, and within-slide spatial artifacts, and the risk of absorbing biologically meaningful spatial signal during that removal. Most normalization strategies address these issues as a preprocessing step, producing normalized values that are subsequently treated as fixed inputs for downstream statistical models. This separation implicitly ignores the uncertainty introduced during normalization, which can lead to biased effect size estimates and underestimation of uncertainty in downstream inference. In this work we propose GPSNorm, a spatially aware Bayesian normalization framework that jointly models technical variation, spatial structure, and biological effects. By embedding normalization directly within a hierarchical probabilistic model, GPSNorm allows uncertainty from the normalization step to propagate naturally into downstream differential expression inference.

The central conceptual contribution of this work is the formulation of normalization and downstream inference as a single joint modeling problem. Rather than estimating size factors or transformed expression values independently of subsequent analysis, GPSNorm treats the technical and biological components of expression as latent variables within a unified statistical model. This formulation enables posterior inference on normalized expression and differential expression effects while accounting for uncertainty in all model components. As a result, uncertainty from normalization is directly reflected in the posterior distribution of downstream quantities such as log-fold changes and differential expression probabilities. This joint modeling approach is particularly advantageous in spatial transcriptomics datasets where technical and biological sources of variation are often intertwined.

A key advantage of the Bayesian formulation is the ability to share information across genes, spatial units, and slides through the hierarchical structure of the model. Spatial transcriptomics datasets often contain relatively small numbers of spatial units compared to the dimensionality of the gene expression space, particularly in targeted platforms such as NanoString GeoMx DSP. In such settings, purely gene-wise or deterministic normalization procedures may struggle to separate technical variation from biological signal. By contrast, the hierarchical priors in GPSNorm induce shrinkage that stabilizes estimates across genes and spatial locations. This information sharing improves the estimation of null genes and reduces the likelihood of spurious differential expression calls, as observed in the simulation study. At the same time, the model retains sufficient flexibility to recover strong biological signals when present.

Explicit modeling of spatial technical structure also improves inference. Spatial artifacts arising from staining variation, tissue thickness, or imaging effects can introduce smooth technical biases across the tissue that resemble biological gradients. If such structure is not modeled explicitly, normalization procedures may either fail to remove the artifact or inadvertently absorb biological signal. GPSNorm addresses this challenge by modeling spatial technical effects through a slide-specific intrinsic conditional autoregressive (ICAR) field, allowing smooth technical variation to be estimated and removed while preserving biological contrasts. The simulation results demonstrate that this approach can successfully recover the underlying technical spatial field while maintaining accurate estimation of biological log-fold changes.

The empirical results on three real spatial transcriptomics datasets further illustrate the practical advantages of the proposed approach. In the human dorsolateral prefrontal cortex (DLPFC) Visium dataset, GPSNorm recovers the expected enrichment of the oligodendrocyte marker gene *MOBP* within white matter regions despite substantial library size heterogeneity across spots. Previous work introducing SpaNorm demonstrated that normalization methods that ignore spatial coordinates often recover enrichment only at the boundary of white matter regions rather than within the region itself. Similar to SpaNorm, GPSNorm is able to recover both boundary and within-region enrichment. However, because GPSNorm is embedded in a Bayesian framework, it additionally provides posterior uncertainty estimates for the inferred spatial expression surface, allowing the strength of enrichment to be quantified probabilistically.

The GeoMx COVID-19 lung damage dataset provides an example of a challenging small-sample spatial transcriptomics study. With only 46 spatial regions distributed across multiple slides, many normalization methods struggle to produce stable differential expression estimates. In this setting, the hierarchical structure of GPSNorm allows the model to borrow strength across genes and spatial units, improving the stability of inference. Moreover, the GeoMx platform provides experimentally defined housekeeping and negative control probes, which are explicitly incorporated into the GPSNorm model to help distinguish technical background variation from biological signal. Using this framework, GPSNorm identifies chemokine genes such as *CCL18, CXCL10*, and *CCL2* as up-regulated in severely damaged lung regions, consistent with the inflammatory signatures reported in the original study.

The Spatial Organ Atlas kidney dataset further highlights the ability of GPSNorm to preserve biologically meaningful transcriptional contrasts. Using curated marker gene sets for glomerular and proximal tubule compartments, we evaluated how well different normalization methods prioritize biologically expected signals. GPSNorm maintained high ranking of glomerular markers while simultaneously prioritizing of proximal tubule markers when comparing glomerular versus non-glomerular regions. This balanced prioritization across compartments suggests that GPSNorm can improve both statistical prioritization of marker genes and biological interpretability of differential expression results.

Despite these promising results, several limitations remain. First, although INLA enables efficient approximate inference for latent Gaussian models, the computational cost of fitting hierarchical spatial models can still be substantial for very large spatial transcriptomics datasets. In practice, this challenge can be mitigated through strategies such as clustering spatial units prior to model fitting, though further work is needed to develop scalable approximations for increasingly large datasets. Second, the current model assumes a particular decomposition of technical and biological variation and relies on smooth spatial priors for technical effects. While this assumption is reasonable for many datasets, more complex technical artifacts or highly irregular spatial patterns may require alternative spatial priors or model extensions.

Future work may extend this framework in several directions. One promising direction is the joint modeling of normalization, differential expression, and spatially variable gene detection within a unified probabilistic framework. Another avenue involves integrating single-cell reference data to improve biological interpretation of spatial signals. Finally, scalable approximations to spatial Gaussian processes may allow similar modeling strategies to be applied to emerging high-resolution spatial transcriptomics technologies that generate orders of magnitude more spatial units.

In summary, GPSNorm demonstrates that treating normalization as a latent statistical inference problem rather than a deterministic preprocessing step can substantially improve downstream analysis in spatial transcriptomics studies. By jointly modeling technical variation, spatial structure, and biological signal within a Bayesian hierarchical framework, GPSNorm provides an approach for propagating normalization uncertainty into differential expression inference while preserving biologically meaningful spatial patterns.

## Data Availability Statement

The 10x Visium Human DLPFC Dataset was obtained through the [SpaNorm R package](https://bioconductor.org/packages//release/bioc/vignettes/SpaNorm/inst/doc/SpaNorm.html). The GeoMx COVID-19 Lung Damage Dataset can be found at Gene Expression Omnibus series GSE186213. The GeoMx Spatial Organ Atlas Kidney Dataset is available at https://brukerspatialbiology.com/products/geomx-digital-spatial-profiler/spatial-organ-atlas/human-kidney/. All preprocessing and analysis code is available at https://github.com/Tiny-Quant/GPSNorm.

## REFERENCES

Julian Besag. Spatial Interaction and the Statistical Analysis of Lattice Systems. 36(2):192–225. ISSN 1369-7412, 1467-9868. doi: 10.1111/j.2517-6161.1974.tb00999.x. URL https://academic.oup.com/jrsssb/article/36/2/192/7027394.

Julian Besag, Jeremy York, and Annie Mollié. Bayesian image restoration, with two applications in spatial statistics. 43(1): 1–20. ISSN 1572-9052. doi: 10.1007/BF00116466. URL https://doi.org/10.1007/BF00116466.

Dharmesh D. Bhuva, Chin Wee Tan, Agus Salim, Claire Marceaux, Marie A. Pickering, Jinjin Chen, Malvika Kharbanda, Xinyi Jin, Ning Liu, Kristen Feher, Givanna Putri, Wayne D. Tilley, Theresa E. Hickey, Marie-Liesse Asselin-Labat, Belinda Phipson, and Melissa J. Davis. Library size confounds biology in spatial transcriptomics data. 25(1):99. ISSN 1474-760X. doi: 10.1186/s13059-024-03241-7. URL https://doi.org/10.1186/s13059-024-03241-7.

Tjessa Bondue, Fanny O. Arcolino, Koenraad R. P. Veys, Oyindamola C. Adebayo, Elena Levtchenko, prefix=van den useprefix=true family=Heuvel, given=Lambertus P., and Mohamed A. Elmonem. Urine-Derived Epithelial Cells as Models for Genetic Kidney Diseases. 10(6): 1413. ISSN 2073-4409. doi: 10.3390/cells10061413. URL https://www.mdpi.com/2073-4409/10/6/1413.

Amy R. Cross, prefix=de useprefix=true family=Andrea, given=Carlos E. María Villalba-Esparza, Manuel F. Landecho, Lucia Cerundolo, Praveen Weeratunga, Rachel E. Etherington, Laura Denney, Graham Ogg, Ling-Pei Ho, Ian S.D. Roberts, Joanna Hester, Paul Klenerman, Ignacio Melero, Stephen N. Sansom, and Fadi Issa. Spatial transcriptomic characterization of COVID-19 pneumonitis identifies immune circuits related to tissue injury. 8(2):e157837. ISSN 2379-3708. doi: 10.1172/jci.insight.157837. URL https://www.ncbi.nlm.nih.gov/pmc/articles/PMC9977306/.

Ciaran Evans, Johanna Hardin, and Daniel M Stoebel. Selecting between-sample RNA-Seq normalization methods from the perspective of their assumptions. 19(5):776–792. ISSN 1477-4054. doi: 10.1093/bib/bbx008. URL https://doi.org/10.1093/bib/bbx008.

Christoph Hafemeister and Rahul Satija. Normalization and variance stabilization of single-cell RNA-seq data using regularized negative binomial regression. 20:296. ISSN 1474-7596. doi: 10.1186/s13059-019-1874-1. URL https://pmc.ncbi.nlm.nih.gov/articles/PMC6927181/.

Laleh Haghverdi, Aaron T. L. Lun, Michael D. Morgan, and John C. Marioni. Batch effects in single-cell RNA sequencing data are corrected by matching mutual nearest neighbours. 36 (5):421–427. ISSN 1087-0156. doi: 10.1038/nbt.4091. URL https://pmc.ncbi.nlm.nih.gov/articles/PMC6152897/.

James S. Hodges and Brian J. Reich. Adding Spatially-Correlated Errors Can Mess Up the Fixed Effect You Love. 64(4):325–334. ISSN 0003-1305. doi: 10.1198/tast.2010.10052. URL https://doi.org/10.1198/tast.2010.10052.

Xiaoqi Liang, Yue Cao, and Jean Yee Hwa Yang. Multi-task benchmarking of spatially resolved gene expression simulation models. URL https://www.biorxiv.org/content/10.1101/2024.05.29.596418v1.

Finn Lindgren and Håvard Rue. Bayesian Spatial Modelling with R-INLA. 63:1–25. ISSN 1548-7660. doi: 10.18637/jss.v063.i19. URL https://doi.org/10.18637/jss.v063.i19.

Finn Lindgren, Håvard Rue, and Johan Lindström. An explicit link between Gaussian fields and Gaussian Markov random fields: The stochastic partial differential equation approach. 73(4):423–498. ISSN 1467-9868. doi: 10.1111/j.1467-9868.2011.00777.x. URL https://onlinelibrary.wiley.com/doi/abs/10.1111/j.1467-9868.2011.00777.x.

Ning Liu, Dharmesh D Bhuva, Ahmed Mohamed, Micah Bokelund, Arutha Kulasinghe, Chin Wee Tan, and Melissa J Davis. standR: Spatial transcriptomic analysis for GeoMx DSP data. 52(1):e2, a. ISSN 0305-1048. doi: 10.1093/nar/gkad1026. URL https://doi.org/10.1093/nar/gkad1026.

Qi Liu, Liang Yue, Jiu Deng, Yingxia Tan, and Chengjun Wu. Progress and breakthroughs in human kidney organoid research. 39:101736, b. ISSN 2405-5808. doi: 10.1016/j.bbrep.2024.101736. URL https://pmc.ncbi.nlm.nih.gov/articles/PMC11190488/.

Michael I Love, Wolfgang Huber, and Simon Anders. Moderated estimation of fold change and dispersion for RNA-seq data with DESeq2. 15(12):550. ISSN 1474-7596. doi: 10.1186/s13059-014-0550-8. URL https://pmc.ncbi.nlm.nih.gov/articles/PMC4302049/.

David Lähnemann, Johannes Köster, Ewa Szczurek, Davis J. McCarthy, Stephanie C. Hicks, Mark D. Robinson, Catalina A. Vallejos, Kieran R. Campbell, Niko Beerenwinkel, Ahmed Mahfouz, Luca Pinello, Pavel Skums, Alexandros Stamatakis, Camille Stephan-Otto Attolini, Samuel Aparicio, Jasmijn Baaijens, Marleen Balvert, prefix=de-useprefix=false family=Barbanson, given=Buys, Antonio Cappuccio, Giacomo Corleone, Bas E. Dutilh, Maria Florescu, Victor Guryev, Rens Holmer, Katharina Jahn, Thamar Jessurun Lobo, Emma M. Keizer, Indu Khatri, Szymon M. Kielbasa, Jan O. Korbel, Alexey M. Kozlov, Tzu-Hao Kuo, Boudewijn P.F. Lelieveldt, Ion I. Mandoiu, John C. Marioni, Tobias Marschall, Felix Mölder, Amir Niknejad, Alicja Raczkowska, Marcel Reinders, prefix=de-useprefix=false family=Ridder, given=Jeroen, Antoine-Emmanuel Saliba, Antonios Somarakis, Oliver Stegle, Fabian J. Theis, Huan Yang, Alex Zelikovsky, Alice C. McHardy, Benjamin J. Raphael, Sohrab P. Shah, and Alexander Schönhuth. Eleven grand challenges in single-cell data science. 21: 31. ISSN 1474-7596. doi: 10.1186/s13059-020-1926-6. URL https://pmc.ncbi.nlm.nih.gov/articles/PMC7007675/.

Kristen R. Maynard, Leonardo Collado-Torres, Lukas M. Weber, Cedric Uytingco, Brianna K. Barry, Stephen R. Williams, Joseph L. Catallini, Matthew N. Tran, Zachary Besich, Madhavi Tippani, Jennifer Chew, Yifeng Yin, Joel E. Kleinman, Thomas M. Hyde, Nikhil Rao, Stephanie C. Hicks, Keri Martinowich, and Andrew E. Jaffe. Transcriptome-scale spatial gene expression in the human dorsolateral prefrontal cortex. 24(3): 425–436. ISSN 1546-1726. doi: 10.1038/s41593-020-00787-0. URL https://www.nature.com/articles/s41593-020-00787-0.

Nghia Millard, Jonathan H. Chen, Mukta G. Palshikar, Karin Pelka, Maxwell Spurrell, Colles Price, Jiang He, Nir Hacohen, Soumya Raychaudhuri, and Ilya Korsunsky. Batch correcting single-cell spatial transcriptomics count data with Crescendo improves visualization and detection of spatial gene patterns. 26:36. ISSN 1474-7596. doi: 10.1186/s13059-025-03479-9. URL https://pmc.ncbi.nlm.nih.gov/articles/PMC11863647/.

NanoString Technologies. Nanostring geomx spatial organ atlas: Human kidney (whole transcriptome atlas). Online dataset, 2026. Available at: https://nanostring.com/products/geomx-digital-spatial-profiler/spatial-organ-atlas/human-kidney/.

Brenda Pardo, Abby Spangler, Lukas M. Weber, Stephanie C. Hicks, Andrew E. Jaffe, Keri Martinowich, Kristen R. Maynard, and Leonardo Collado-Torres. spatiallibd: an r/bioconductor package to visualize spatially-resolved transcriptomics data. BMC Genomics, 2022. doi: 10.1186/s12864-022-08601-w. URL https://doi.org/10.1186/s12864-022-08601-w.

Jingyang Qian, Hudong Bao, Xin Shao, Yin Fang, Jie Liao, Zhuo Chen, Chengyu Li, Wenbo Guo, Yining Hu, Anyao Li, Yue Yao, Xiaohui Fan, and Yiyu Cheng. Simulating multiple variability in spatially resolved transcriptomics with scCube. 15(1):5021. ISSN 2041-1723. doi: 10.1038/s41467-024-49445-0. URL https://www.nature.com/articles/s41467-024-49445-0.

Carl Edward Rasmussen and Christopher K. I. Williams. Gaussian Processes for Machine Learning. Adaptive Computation and Machine Learning. MIT Press, 3. print edition. ISBN 978-0-262-18253-9.

Brian J. Reich, James S. Hodges, and Vesna Zadnik. Effects of Residual Smoothing on the Posterior of the Fixed Effects in Disease-Mapping Models. 62(4):1197–1206. ISSN 0006-341X. doi: 10.1111/j.1541-0420.2006.00617.x. URL https://doi.org/10.1111/j.1541-0420.2006.00617.x.

Davide Risso, John Ngai, Terence P. Speed, and Sandrine Dudoit. Normalization of RNA-seq data using factor analysis of control genes or samples. 32(9):896–902. ISSN 1546-1696. doi: 10.1038/nbt.2931. URL https://www.nature.com/articles/nbt.2931.

Mark D Robinson and Alicia Oshlack. A scaling normalization method for differential expression analysis of RNA-seq data. 11 (3):R25. ISSN 1474-7596. doi: 10.1186/gb-2010-11-3-r25. URL https://pmc.ncbi.nlm.nih.gov/articles/PMC2864565/.

Havard Rue and Leonhard Held. Gaussian Markov Random Fields: Theory and Applications. Chapman and Hall/CRC. ISBN 978-0-429-20882-9. doi: 10.1201/9780203492024.

Håvard Rue, Sara Martino, and Nicolas Chopin. Approximate Bayesian inference for latent Gaussian models by using integrated nested Laplace approximations. 71(2):319–392. ISSN 1467-9868. doi: 10.1111/j.1467-9868.2008.00700.x. URL https://onlinelibrary.wiley.com/doi/abs/10.1111/j.1467-9868.2008.00700.x.

Agus Salim, Dharmesh D. Bhuva, Carissa Chen, Chin Wee Tan, Pengyi Yang, Melissa J. Davis, and Jean Y. H. Yang. SpaNorm: Spatially-aware normalization for spatial transcriptomics data. 26(1):109. ISSN 1474-760X. doi: 10.1186/s13059-025-03565-y. URL https://doi.org/10.1186/s13059-025-03565-y.

Daniel Simpson, Håvard Rue, Andrea Riebler, Thiago G. Martins, and Sigrunn H. Sørbye. Penalising Model Component Complexity: A Principled, Practical Approach to Constructing Priors. 32(1). ISSN 0883-4237. doi: 10.1214/16-STS576. URL https://projecteuclid.org/journals/statistical-science/volume-32/issue-1/Penalising-Model-Component-Complexity--A-Principled-Practical-Approa10.1214/16-STS576.full.

Patrik L. Ståhl, Fredrik Salmén, Sanja Vickovic, Anna Lundmark, José Fernández Navarro, Jens Magnusson, Stefania Giacomello, Michaela Asp, Jakub O. Westholm, Mikael Huss, Annelie Mollbrink, Sten Linnarsson, Simone Codeluppi, Åke Borg, Fredrik Pontén, Paul Igor Costea, Pelin Sahlén, Jan Mulder, Olaf Bergmann, Joakim Lundeberg, and Jonas Frisén. Visualization and analysis of gene expression in tissue sections by spatial transcriptomics. 353(6294):78–82. ISSN 1095-9203. doi: 10.1126/science.aaf2403.

